# IDHwt glioblastomas can be stratified by their transcriptional response to standard treatment, with implications for targeted therapy

**DOI:** 10.1101/2023.02.03.526945

**Authors:** Georgette Tanner, Rhiannon Barrow, Martina Finetti, Shoaib Ajaib, Nazia Ahmed, Steven Pollock, Nora Rippaus, Alexander F. Bruns, Khaja Syed, James Poulter, Erica Wilson, Colin Johnson, Frederick S. Varn, Anke Brüning-Richardson, Catherine Hogg, Alastair Droop, Arief Gusnanto, Matthew A. Care, Luisa Cutillo, David Westhead, Susan C. Short, Michael D. Jenkinson, Andrew Brodbelt, Aruna Chakrabarty, Azzam Ismail, Roel GW Verhaak, Lucy F. Stead

**Affiliations:** Leeds Institute of Medical Research at St James’s, University of Leeds, Leeds, UK; Leeds Institute of Cardiovascular and Metabolic Medicine, University of Leeds, Leeds, UK; The Jackson Laboratory for Genomic Medicine, Farmington, CT, USA; School of Applied Sciences, University of Huddersfield, Huddersfield UK; Welcome Sanger Institute, Hinxton, Saffron Walden, UK; School of Mathematics, University of Leeds, Leeds, UK; The Walton Centre NHS Foundation Trust, Liverpool, UK; Leeds Teaching Hospital, Leeds, England; Institute of Systems, Molecular and Integrative Biology, University of Liverpool, Liverpool, UK

## Abstract

Glioblastoma (GBM) brain tumours lacking *IDH1* mutations (IDHwt) have the worst prognosis of all brain neoplasms. Patients receive surgery and chemoradiotherapy but tumours almost always fatally recur. Using RNAseq data from 107 pairs of pre- and post-standard treatment locally recurrent IDHwt GBM tumours, we identified two responder subtypes based on therapy-driven changes in gene expression. In two thirds of patients a specific subset of genes is up-regulated from primary to recurrence (Up responders) and in one third the same genes are down-regulated (Down responders). Characterisation of the responder subtypes indicates subtype-specific adaptive treatment resistance mechanisms. In Up responders treatment enriches for quiescent proneural GBM stem cells and differentiated neoplastic cells with increased neurotransmitter signalling, whereas Down responders commonly undergo therapy-driven mesenchymal transition. Stratifying GBM tumours by response subtype may lead to more effective treatment. In support of this, modulators of gamma aminobutyric acid (GABA) neurotransmitter signalling differentially sensitise Up and Down responder GBM models to standard treatment *in vitro*.

## Introduction

Glioblastoma (GBM) brain tumours are currently incurable. GBM cancer cells infiltrate the surrounding normal brain so, although patients undergo surgery, complete resection is not possible. Standard of care includes subsequent chemoradiation with Temozolomide (TMZ), but this only modestly prolongs survival: typically tumours recur 6-9 months later and are almost always fatal. In order to kill these cells more effectively we must understand why some unresected GBM cells survive chemo radiation. To this end, global efforts have characterised clonal evolution through treatment in GBM. We performed the largest of such genomic studies and, in agreement with Körber et al., concluded that treatment resistance is not driven by somatic mutations in specific genes or pathways[1, 2]. Single cell analyses of primary GBM tumours have identified transcriptionally-defined neoplastic cell states that are shared across genomic subclones and patients, and exhibit high levels of plasticity[3–5]. Single cell epigenetic profiling of primary tumours concurs with this notion, having linked chromatin accessibility with the observed neoplastic transcriptional states[6, 7].Together, these findings suggest that epigenetics underpins GBM cell behaviours, including those that enable inherent or adaptive treatment resistance.

Our subsequent efforts have focused on characterising the changes in transcriptional profiles through treatment. Our initial investigation of bulk longitudinal glioma profiles indicated a convergence upon specific phenotypes at recurrence[8]. However, distinctly different trajectories occur for tumours that are wild-type for isocitrate dehydrogenases (IDHwt) compared to those harbouring mutations in the genes encoding these enzymes (IDHmut)[8], indicating that these two entities require separate analysis[8]. Wang et al. previously showed that primary GBM cancer cells reside on a single axis of variation between proneural and mesenchymal phenotypes[5]. This group have more recently performed multi-omic, single cell analyses of longitudinal IDHwt GBM tumours and shown that this is also true for recurrent tumour but that treatment alters the prevalence of cells along this axis, whilst highlighting mechanisms by which a mesenchymal shift may occur[9].

This study described herein expands upon previous bulk tumour studies by performing transcriptional analysis of the largest cohort (n=214) of paired longitudinal GBMs that are specifically IDHwt and recurred locally following standard treatment. We identify a subset of genes, linked by an epigenetic remodelling complex, that are consistently dysregulated through standard of care but in opposite directions in different patients. This defines two responder subtypes: ‘Up’ and ‘Down’ responders. We show that this stratification of IDHwt GBM patients is biologically and clinically meaningful, suggesting responder subtype-specific treatment resistance mechanisms and therapeutic vulnerabilities. To investigate this further we identified *in vitro* models that recapitulate the responder subtypes. We then pharmaceutically modulated a key aspect of differential responder biology, gamma-aminobutyric acid (GABA) signalling, and observed the effect on cell survival following administration of radiation and TMZ. We found opposing impacts on the efficacy of non-surgical components of standard of care in the responder subtype models, suggesting that stratified adjuvant therapies are required to combat treatment resistance in IDHwt GBM. Our work confirms our previous findings in longitudinal bulk samples, and those of Wang et al. at single cell resolution, whilst providing additional context that increases the translational impact.

## Results

### Overview of Discovery and Validation cohorts

We restricted our study to *de novo* IDH^wt^ GBM tumours and matched, post-standard treatment (having received radiation and Temozolomide), locally recurrent tumours. The Discovery cohort consisted of 168 longitudinally paired samples from 84 patients for which RNAseq data was processed in house. The Validation cohort consisted of 46 paired samples from 23 patients for which RNAseq data was processed via a distinct pipeline within the Glioma Longitudinal AnalysSiS (GLASS) consortium[8] (**Fig. 1A; Table S1**). Sequencing metrics for the Discovery cohort are given in **Table S2**.

**Figure 1.**
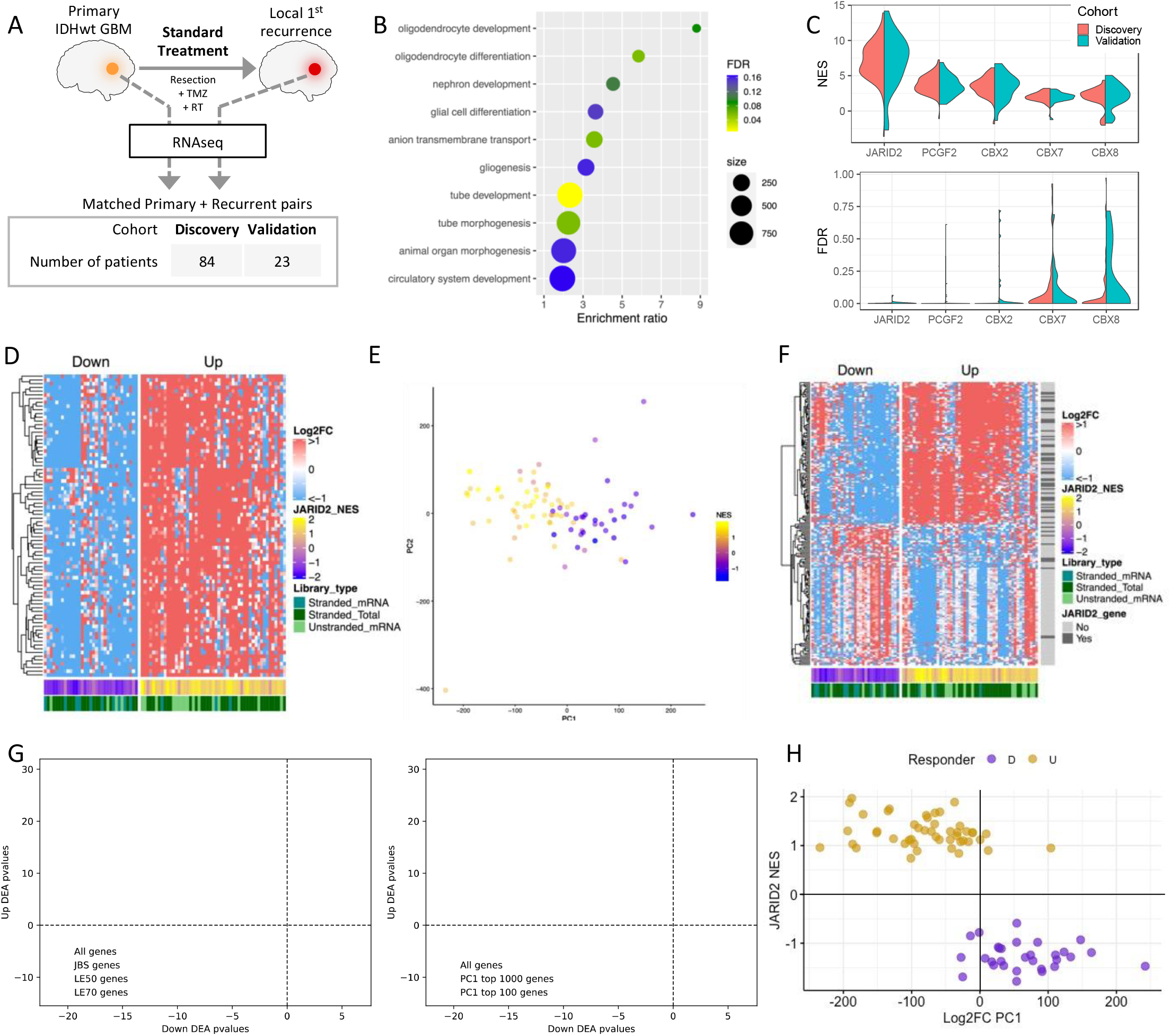
**A**. A schematic outlining the study design and cohort sizes. The figures in this panel visualise data from the Discovery cohort (Validation cohort data is displayed in supplemental figures). **B**. The biological processes enriched in the genes differentially expressed between matched primary and recurrent GBMs. **C**. The distribution of per-patient normalised enrichment scores (NES, top plot) and false Discovery rates (FDR, bottom plot) for the top-scoring promoter-binding factors associated with gene expression changes between recurrent vs primary GBMs. **D**. A heatmap of the fold change in expression between recurrent vs primary GBMs samples for each patient (columns) for the JARID2 gene site binding sites genes (JBSgenes) that were in the leading edge of the GSEA results in more than 50% of patients (LE50 genes, rows). Patients separate into Up (JARID2 gene set NES>0) and Down (NES<0) responders irrespective of the heterogeneity in RNAseq library preparation approach. **E**. Patients are plotted, coloured by JBSgenes NES, according to principal components 1 (PC1) and 2 (PC2) of their whole transcriptome fold change in expression between their recurrent vs primary GBM. **F**. A heatmap of the fold change in expression between recurrent vs primary GBMs samples for each patient (columns) for the largest 100 positive and largest 100 negative weighted genes of PC1 from panel E (rows). Whether each gene is a JBSgene is also indicated. **G**. Each gene is plotted according to its −log_10_p-value result of separate differential gene expression analyses in matched recurrent vs primary tumours in Down (x-axis) and Up (y-axis) responders. Left plot: genes coloured according to whether they are JBSgenes or, more specifically, LE50 and LE70 genes. Right plot: genes coloured according to whether they are in the top 100 or 1000 genes ranked by absolute value of PC1 ranking from the analysis in panel E. **H**. Plotting patients according to their JBSgene NES and PC1 score from panel E clearly separates of Up and Down responders.

### Therapy-driven changes in gene expression delineate two responder subtypes in GBM

The genes significantly differentially expressed between matched primary and recurrent GBMs were enriched for terms associated with neurodevelopment (**Fig. 1B**). Neurodevelopment is orchestrated by master regulators of gene expression, where cascades of transcription factors work in concert with chromatin remodelling complexes. To investigate whether specific regulators were implicated in the genes dysregulated in primary versus recurrent GBMs, we created novel, comprehensive gene sets for DNA-binding factors. Using these in gene set enrichment analysis, we found that genes containing a JARID2 (Jumonji and AT-Rich Interacting Domain 2) binding site in their promoter (JBSgenes) were consistently and significantly altered through treatment across patients in both the Discovery and Validation cohorts (**Fig. 1C; Tables S3-12**). This result was consistent when the definition of a promoter was extended from 1kb to 2kb or 5kb either side of the transcription start site (TSS) and when quantifying expression at the level of individual TSSs, rather than genes (**Fig. S1A and B; Tables S13-18**).

To assess whether the same JBSgenes were driving the enrichment across patients we inspected the stability of inclusion within the leading edge. In a total of 5234 JBSgenes, 443 were LE50 (genes in the leading edge of at least 50% of patients) and 81 were LE70 in the Discovery cohort. These significantly overlapped with the LE50 (444 genes) and LE70 (87genes) calculated independently in the Validation cohort (hypergeometric test P=8.2E-217 and P=6.2E-40 and representation factors of 7.3 and 24.5 respectively) (**Table S19**). The fold change in expression from primary to recurrence (Log2FC) for LE70 genes was in a consistent direction within, but differed between, patients (**Fig. 1D**). In 60% of patients LE70 genes were upregulated from primary to recurrence (coined Up responders), and in the remaining 40% the same genes were downregulated (Down responders). This phenomenon was recapitulated in the Validation cohort with the same split of Up and Down responders (chi-squared test P=0.98; **Fig. S1C**).

JARID2 is an accessory protein to Polycomb Repressive Complex 2 (PRC2), which is responsible for gene repression via deposition of H3K27me3. PRC2 has a role in the transcriptional reprogramming required for neurodevelopment and brain cell lineage determination. To investigate transcriptional reprogramming from primary to recurrent GBM we performed principal component analysis on Log2FC profiles. Principal component 1 (PC1), the main source of variation, separated patients according to the strength of their classification as Up or Down responders, quantified by their JARID2 gene set normalised enrichment score (NES), in both the Discovery (**Fig. 1E, Table S20**) and Validation cohort (**Fig. S1D, Table S21**). The 100 genes with the highest PC1 loadings were significantly enriched for JBSgenes (chi-squared test P=1.8E-5; **Fig. 1F**). This indicates that Up and Down responder tumours undergo transcriptional reprogramming in opposing directions through treatment, driven by a key set of genes but propagated throughout the transcriptome. Splitting patients by responder subtype and performing separate primary versus recurrent differential gene expression analyses further confirmed that the same genes are being dysregulated in different directions, with the most significant being enriched in JBSgenes and those with the highest PC1 loadings (**Fig. 1G; Table S22-23**). Plotting patients according to their Log2FC PC1 value and their JARID2 NES resulted in a clear separation of Up and Down responders (**Fig. 1H**)

### Responder subtypes undergo different changes in neoplastic and immune cell populations through treatment

We proceeded to investigate the clinical and biological differences between responder subtypes. In multivariate survival analysis, including known prognostic GBM markers (i.e. expression of O-6-Methylguanine-DNA Methyltransferase (*MGMT)* in the primary tumour and age at diagnosis) there was no association between responder type and progression free (P=0.59, β=−0.86) or overall (P=0.97, β=0.1) survival. The prevalence of classical, mesenchymal and proneural GBM subtypes in primary tumours is the same for both responder subtypes (chi-squared P=0.98), and the probability of switching subtype between primary and recurrence remains the same (chi-squared P=0.89). However, there is a significant difference in subtype representation between responder subtypes at recurrence (chi-squared P=0.0030 **Fig. 2A**). In Up responders the majority of samples that switch subtype become proneural (60%, n=16) whereas in Down responders there is a significant switch to mesenchymal subtype (61%, n=11). The same is observed in the Validation cohort where 43% of Up responders that switch become proneural (n=7) and 67% of Down responders that switch become mesenchymal (n=6) (**Fig. S2A**). GBM tumour subtypes, defined by bulk RNAseq profile clustering, result from the dominant single GBM cell type signal in the GBM sample at any given timepoint[3]. The prevalence of single GBM cell types is influenced by the tumour microenvironment[3]. To investigate whether the opposing gene dysregulation observed in Up and Down responders resulted from different cell population dynamics, we developed and validated GBMdeconvoluteR (https://gbmdeconvoluter.leeds.ac.uk), a GBM-specific neoplastic and immune cell deconvolution tool, and applied this to our datasets[10] (**Fig. 2B and S2B**). GBMdeconvoluteR uses the neoplastic GBM cell classifications derived in Neftel et al., quantifying the prevalence of mesenchymal (MES), astrocyte cell like (AC), neural progenitor cell like (NPC) and oligodendrocyte progenitor cell like (OPC) cancer cells[3]. We found that neoplastic cell population changes in response to treatment are significantly different between the responder subtypes (**Fig. 2B)**: Down responders predominantly increase in MES GBM cells (Wilcoxon P=1.6E-3) and decrease in both NPC (Wilcoxon P=8.0E-4) and OPC (Wilcoxon P=5.6E-5) cells. In Up responder, the opposite is true, with OPC cells increasing the most through treatment. In both responder subtypes AC cells are more likely to decrease through treatment, but significantly more so in Down responders than Up (Wilcoxon P=0.047). These subtype-specific changes in neoplastic cells are also observed in the Validation dataset (**Fig. S2B**). Conversely changes in immune cell population are less evident through treatment in Up and Down responders and significant findings differ between the Discovery cohort (where NK cell infiltration is significantly increased in Up responders: Wilcoxon P=9.8E-4 **Fig. 2B**) and Validation cohort (where T cell infiltration is significantly increased in Up responders: Wilcoxon P=0.016, and monocyte infiltration is significantly increased in Down responders Wilcoxon P=0.028; **Fig. S2B**). Upon re-classifying cells as lymphoid or myeloid across both datasets we found that Up responders show a significant increase in lymphoid cells through treatment (Wilcoxon P=8.8E-4 **Fig. S2C**).

**Figure 2.**
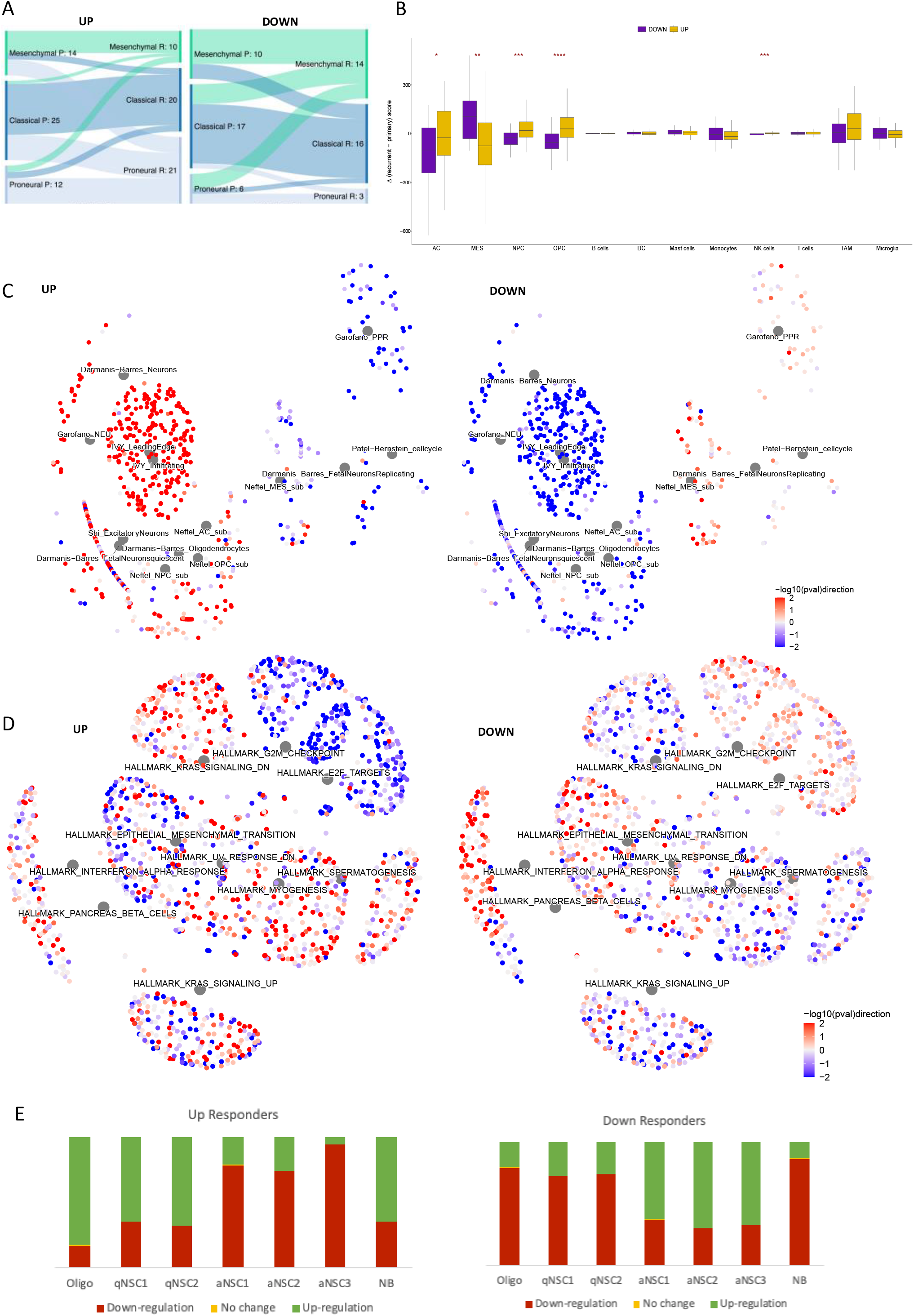
**A**. Sankey plots showing the prevalence of subtype switching from primary (P) to recurrent (R) GBM in the Up responders (left) and Down responders (right) in the Discovery cohort. **B**. The distributions of change in cell type score, assigned per sample by GBMdeconvoluteR to indicate the prevalence of that cell type in the tumour, between primary and matched recurrent GBMs in Down (purple) and Up (gold) responders. A dotted line indicates no change. The median is noted by a black horizontal line. Significance is denoted by asterisks: *: p<0.05; **: p<0.01; ***: p<0.001; ****: p<0.0001. Neoplastic GBM cells are on the left of the plot: AC= astrocyte like; MES= mesenchymal like; NPC= neural progenitor like; OPC= oligodendrocyte progenitor like. **C**. Network plots showing the GBM biology specific gene sets (described in **Table S3**) that are significantly enriched (FDR<0.05) in either, or both, Up or Down responders through treatment. Large grey hub nodes indicate the gene sets. These have associated, smaller, leaf nodes signifying the genes in that set, coloured according to the strength and direction of differential expression through treatment (quantified as −log_10_p-value multiplied by the direction of fold change: −log10(pval)direction) in Up responders (left image) or Down responders (right image). **D**. As for panel C but visualising the Hallmark gene sets from MSigDB that were enriched with an FDR<0.25 in either, or both, responder subtypes. **E**. The proportion of neural stem cell markers of quiescence (qNSC) or active cycling (aNSC), or markers of more differentiated neuroblasts (NB) or oligodendrocytes (Oligo), that were upregulated (green), downregulated (burgundy) or stable (yellow) in Up (left) and Down (right) responders.

### Genes that are differentially expressed through treatment in the responder subtypes suggest different adaptive treatment resistance mechanisms

We next performed gene set enrichment analysis on the genes differentially expressed between primary and recurrent GBMs in the responder subtypes separately, to identify distinct biological processes occurring through treatment. To probe brain and GBM biology more specifically we collated custom gene sets, alongside those from the Molecular Signatures Database (MSigDB), from the literature and public databases[3, 5, 11–33]. These included the latest cell type markers from the BRAIN Initiative Cell Census Network and phenotypic GBM gene signatures from recent single cell studies (**Table S24**). Signatures of neuronal (NEU) neoplastic GBM cells, as defined by a study inspecting distinct pathways activated in single cell clusters[32], were significantly up-regulated in Up responders (90% of genes in this set, Garofano_NEU, were significantly upregulated; FDR=0, n=40; **Fig. 2C and Table S25**). This GBM cell type has elevated levels of neurotransmitter receptors implicated in glioma-neuron interactions, including synapses. In Up responders gamma-aminobutyric acid (GABA) neurotransmitter signalling components, specifically, were significantly upregulated (85% of GABA gated chloride ion channel activity genes were significantly upregulated through treatment in Up responders n=13 FDR=0.03 and down-regulated in Down responders, FRD=0.0053; **Table S26**). Gene signatures of specifically differentiated type of neurons, oligodendrocytes, and the infiltrative leading edge from patient GBM samples were also significantly upregulated in Up responders (FDR<0.05). All of these gene sets were significantly down-regulated in Down responders (**Fig. 2C and Table S25**). This suggests increased neuronal signalling and interactions between cancerous and normal brain cells through treatment in Up responders, specifically, alongside genes that demarcate processes of neuronal and glial cell differentiation.

Signatures of developmental glioma stem cell (GSC) states were also upregulated in Up, and downregulated in Down, responders (Richards_Developmental gene set, n=444, FDR=0.10 and 0.07, respectively; 94-97% of DE genes being unidirectionally dysregulated) and, specifically, signatures of oligodendrocyte progenitor cell-like (OPC) neoplastic cells, in agreement with the cell deconvolution results (Neftel_Cell_2019_OPC gene set, n=49, FDR=0.11 and 0.05, respectively; 93-100% of DE genes being unidirectionally dysregulated **Fig. 2B-C and Table S25**). Wang et al. performed single cell transcriptional analysis of GBM tissues, showing that cells can be transcriptionally assigned along a proneural GSC (pGSC) to mesenchymal GSC (mGSC) axis, with more differentiated malignant cells in the middle[5]. pGSCs uniquely expressed OPC genes. Together with the bulk tumour subtyping results, this suggests that Up responders become enriched for pGSCs through treatment. Conversely, in Down responders there is repeated enrichment of mesenchymal GBM cell signatures (**Fig. 2B-C and Table S25**). Within the MSigDB gene sets, there is also enrichment for genes associated with epithelial to mesenchymal transition. These genes are significantly more down-regulated in Up responders and upregulated in Down responders (adjusted Chi-squared P=1.36E-5; **Fig. 2D and Table S27**). In GBM, a non-epithelial cancer, this hallmark process is more accurately referred to as proneural to mesenchymal transition[34]. Across both the MSigDB and custom gene sets, there was evidence that Down responders upregulate cell cycle genes from primary to recurrent, whereas these are being decreased in Up responders (**Fig. 2C-D and Tables S25-27**). Conversely, in Up responders there is significant upregulation of neural stem cell (NSC) quiescence markers. Codega et al. extracted NSCs from adult mouse brains, used label retention approaches to separate those which were quiescent (qNSCs) from those which were activated (aNSCs), and identified differentially expressed genes[31]. We found that a significant number of qNSCs markers were upregulated in Up responders (73%, n=331; NES=1.2, FDR=0.10; **Table S25**) with most of the same genes downregulated in Down responders (73%, n=327; NES=1.2, FDR=0.07; **Table S25**). Llorens-Bobadilla et al. did scRNAseq of murine adult subventricular zone NSCs, isolated via expression of GLAST and Prom1, and identified 7 unsupervised clusters: two defining qNSCs, three defining aNSCs, one defining neuroblasts (NB) and one defining oligodendrocytes (Oligo; which express low levels of both NSC marker proteins)[30]. We included the genes that delineated these clusters in our custom gene sets (**Table S24**). We found that Up responders up-regulate markers of both qNSCs clusters and the more differentiated NB and Oligo cell types, with concomitant downregulation of genes for all three aNSC clusters, whereas the opposite was evident in the Down responders (**Fig. 2E and Table S25**).

### Gene expression correlation networks highlight different genes influencing therapy-driven transcriptional reprogramming in responder subtypes

To investigate what was driving transcriptional reprogramming in opposing directions through treatment we performed parsimonious gene correlation network analysis (PGCNA), creating networks based on the fold change in expression from primary to recurrence (Log_2_FC) in Up and Down responders separately. We calculated the integrated value of influence (IVI) for each gene in each network. IVI summarises numerous network parameters, such as hubness and degree centrality, to provide a metric of the overall importance of any one gene in a network. We investigated the distribution of IVI among non-JBS, JBS, LE50 and LE70 genes and found that the leading-edge genes (LE50 and, even more so, LE70) are significantly more influential on gene expression dysregulation in response to treatment (**Fig. 3A**). We then built separate gene expression networks for Up responder primary samples (Primary Up), and for Primary Down, Recurrent Up and Recurrent Down. LE50 and LE70 genes become significantly more influential with regards to gene expression regulation after treatment in Up responders and significantly less influential in Down responders after treatment (**Fig. 3B)**. These changes are observed across several local and global parameters that quantify gene regulation within each the network: spreading, hubness, betweenness and degree (**Fig. S3**). This implicates the most commonly dysregulated JBS genes, specifically, in driving the whole transcriptome reprograming observed through treatment, and further validates that this reprogramming occurs in opposing directions.

**Figure 3.**
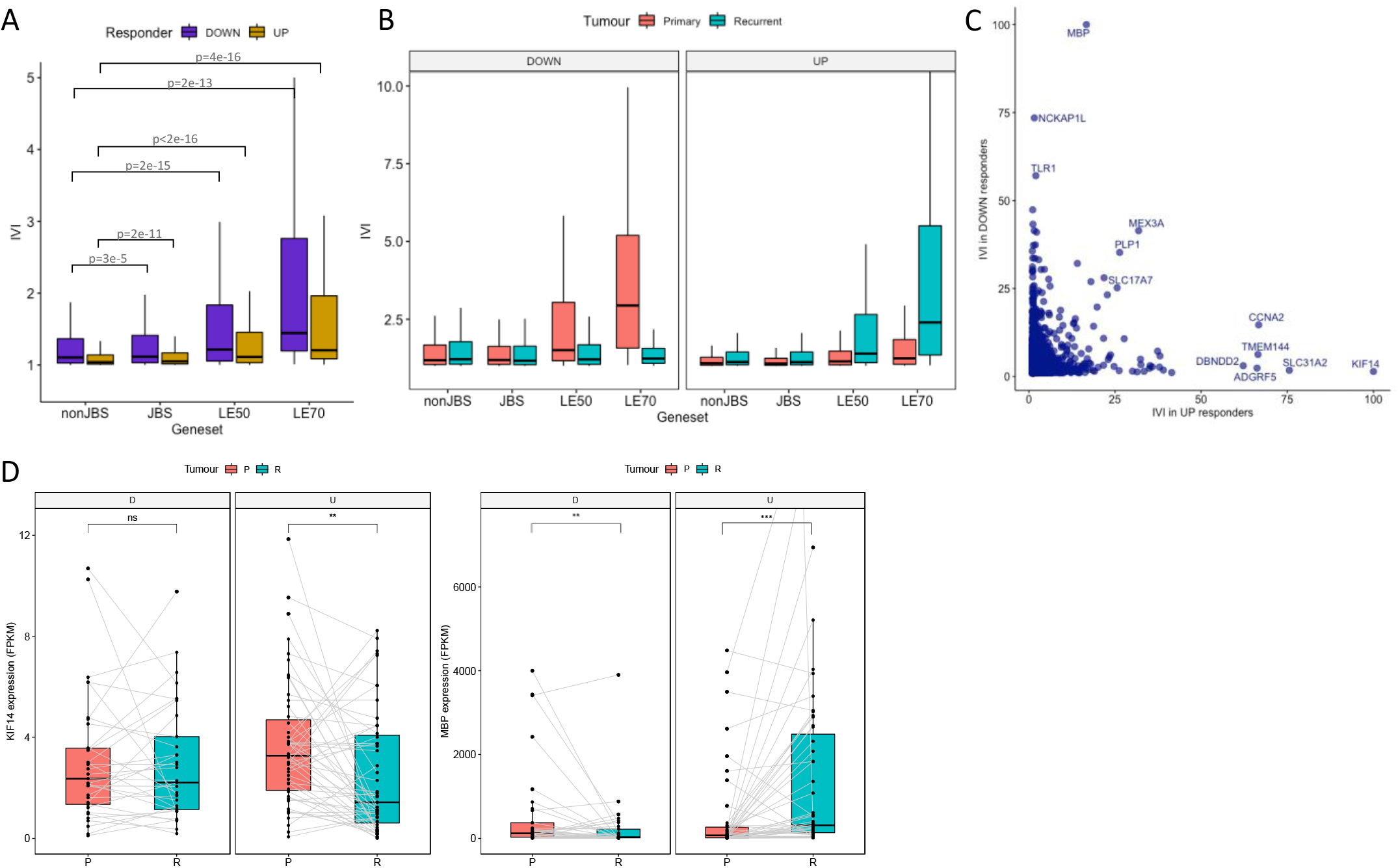
**A**. The distribution of integrated value of influence (IVI) scores for different gene sets, calculated from log2FC (fold change in expression from recurrent to primary) correlation networks, for Down (purple) and Up (gold) responders. nonJBS: genes not in the JARID2 gene set; JBS: genes in the JARID2 geneset but excluding those in the leading edge of at least 50% of patients (LE50 genes); LE50: genes in the LE50 geneset but excluding those in the LE70 geneset; LE70: genes in the leading edge of at least 70% of patients. **B**. As panel A except correlation networks were built from gene expression data in primary (pink) or recurrent (teal) tumours in Down (left panel) or Up (right panel) responders, separately. **C**. Genes are plotted according to their IVI score in log2FC networks of Down (x-axis) and Up (y-axis) responders. Genes that are high in both, or uniquely high in one, log2FC network are labelled. **D**. The expression values for genes encoding Kinesin Family Member 14 (KIF14: left image) and Myelin Basic Protein (MBP: right image) are shown in the primary (pink) and recurrent (teal) tumours of Down (D) and Up (U) responders. Gray lines indicate expression values in primary and recurrent GBMs form the same patient. Significance is denoted: ns: not significant; *: p<0.05; **: p<0.01; ***: p<0.001; ****: p<0.0001.

Comparing individual IVI values in the Up vs Down responder Log_2_FC networks highlighted genes that were uniquely important in coordinating transcriptional reprogramming for each subtype (**Fig. 3C**). Kinesin Family Member 14 (*KIF14*) is highly influential in the Up responders only (IVI is 100 in Up and 1.4 in Down responders, **Fig. 3C**). This gene does not change expression between paired primary and recurrent samples in Down responders but significantly decreases in expression in Up responders (adjusted P=2.0E-3, **Fig. 3D)**. KIF14 is a microtubule motor protein with a role in cell proliferation. Myelin Basic Protein (*MBP*) is highly influential in the Down responders only (IVI is 100 in Down and 16.7 in Up responders, **Fig. 3C**). *MBP* is significantly downregulated through treatment in Down responders and also significantly upregulated in Up responders (adjusted P=8.9E-3 and 2.70E-10 respectively **Fig. 3D**). MBP is a major constituent of the myelin sheath of oligodendrocytes.

This suggests that the genes that are uniquely influential in therapy-driven transcriptional reprogramming in each responder subtype are so because they become down-regulated. Their influence may, therefore, be exerted by becoming rate limiting to the specific processes they associate with, which we showed above to be associated with the opposing responder subtype. This implies that reducing expression of key genes identified through these network analyses could prohibit certain resistance mechanisms.

### Responder subtypes are recapitulated in preclinical models of GBM

Our findings indicate that GBM tumours can be stratified by their transcriptional response to standard treatment, implicating subtype-specific mechanisms of treatment resistance that could yield subtype-specific therapeutic targets. To test this requires experimental models that recapitulate responder subtypes. We cultured an established (A172) and a patient-derived, serum-free (GBM63) GBM cell line as 3D spheroids and subjected them to non-surgical elements of standard treatment: 2Gy irradiation and 30μM TMZ. Treatment caused a significant reduction in both spheroid size (average spheroid size reduced in treated samples by 17.45%: t-test, P=5.2E-3 for A172 and by 18.11%: t-test, P=1.0E-4 for GBM63, n=3) and cell viability (average cell viability reduced in treated samples by 29.46%: t-test, P=0.021 for A172 and by 12.80%: t-test, P=7.0E-4 for GBM63, n=3) in both cell lines in comparison with untreated controls. At the end point, RNA was extracted for each experiment in triplicate. Plotting the fold change in expression between treated and untreated samples (Log2FC) according to the JARID2 normalised enrichment score (NES) and the first principal component (PC1) separated these cell lines as it did for patient primary and recurrent pairs, albeit with smaller effect size. This denoted A172 as an Up, and GBM63 as a Down, responder (**Fig. 4A**). Differential gene expression corroborated the different direction of transcriptional dysregulation in these cell lines in response to treatment: 80% of genes DE in both cell lines were dysregulated in opposite directions (**Fig. 4B**).

**Figure 4.**
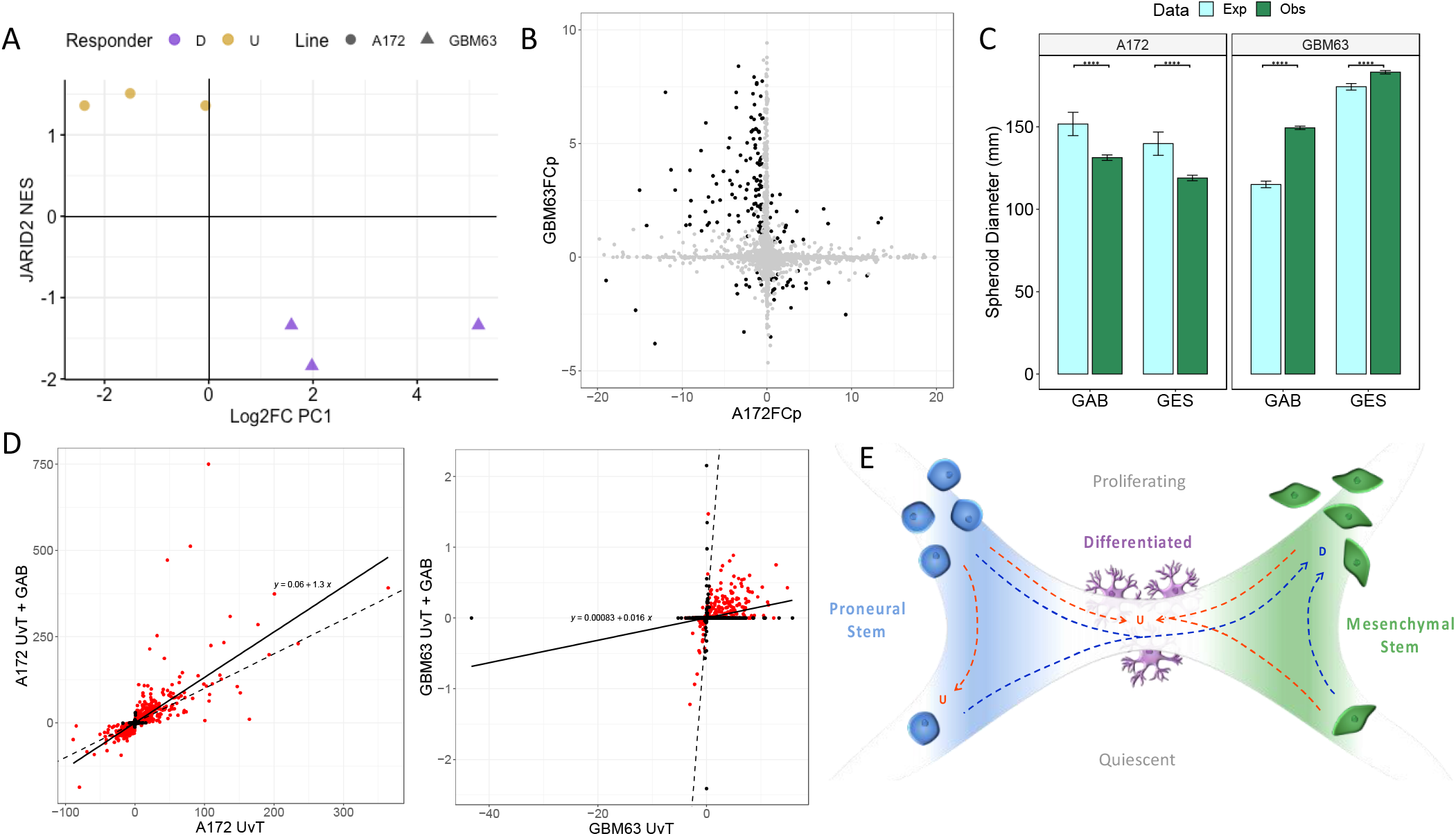
**A**. Replicate experiments in A172 and GBM63 cell line spheroids with and without chemoradiation are plotted according to the JARID2 gene set NES and the value of first principal component when results are projected on the patient principal components shown in Figure 1E. This denotes A172 as an Up responder, and GBM63 as a Down responder, model in correspondence with patient samples as similarly plotted in Figure 1H. **B**. Results of differential expression (DE) analysis between treated and untreated spheroids of A172 and GBM63 separately (n=3). Genes are plotted according to their −log_10_adjusted-p-value multiplied by the log_2_fold change (FCp) in A172 (x-axis) and GBM63 (y-axis) experiments. Black=significantly DE (FDR<0.05) in both cell lines. **C**. The expected (Exp) size of spheroids following administration of chemoradiation and GABA agonist Gaboxadol (GAB) or Guanidinoethyl sulphonate (GES) if there was no interaction between the two (light blue) compared with the observed size (green) in A172 (left) and GBM63 (right). Significance is denoted with asterisks: *: p<0.05; **: p<0.01; ***: p<0.001; ****: p<0.0001. **D**. Genes are plotted according to the size and direction of dysregulation through treatment (the −log_10_adjusted-p-value multiplied by the log_2_fold change between treated and untreated spheroids) without (x-axis) and with (y-axis) administration of the GABA agonist Gaboxadol, for A172 (left) and GBM63 (right) separately (n=3). Red=significantly DE (FDR<0.05) in both conditions. The dotted line is y=x and the solid line indicates the true relation with the equation given on the plot. **E**. Our working model to explain GBM responder subtypes is that GBM cells are on a phenotypic axis between proneural and mesenchymal stem cells as suggested by Wang et al. These stem cells can be in a quiescent or actively cycling state. Differentiated, interconnected (with both each other and surrounding normal cells) cell states lie in the centre of the axis. In Down responders, cells in the GBM tumour move towards the mesenchymal phenotype and increase proliferation rates in response to standard of care. In Up responders, neoplastic GBM cells either become more differentiated and integrate with surrounding cells, upregulating neurotransmitter signalling as they do, or they convert to or remain as proneural stem cells but in a quiescent state in response to standard of care.

### Responder subtype models show differential sensitisation to standard treatment upon the addition of GABA signalling modulators

Gamma aminobutyric acid (GABA) signalling is altered in different directions in Up versus Down responders through treatment (**Table S25)**. We used A172 and GBM63 to inspect how the response to standard treatment changed in the presence of modulators of GABA signalling in Up versus Down responders. We selected two GABA A receptor (GABA_A_R) agonists: guanidinoethyl sulphonate (GES), a weak agonist, and Gaboxadol (GAB), a strong agonist[35, 36]. Linear mixed effects models were built for each cell line independently and showed a significant interaction between the agonist and standard treatment, with regards impact on spheroid size, with P=0 in both cases. However, the modulators altered the response to standard treatment in different directions, with the effect size being proportionate to the strength of agonism (**Fig. 4C**). In A172 the modulator increased the cytotoxic effect of standard treatment (spheroids were 13% smaller on average), whereas in GBM63 it reduced it (spheroids were 10% larger on average). RNA sequencing was performed in triplicate with and without the application of standard treatment in the presence and absence of GAB, the strongest agonist. There was a strong, significant correlation in therapy-driven gene expression changes between GAB and the control for both cell lines (A172: Spearman’s rho=0.81, P=0; GBM63: Spearman’s rho=0.64, P=0). However, in A172 the gene expression changes were significantly amplified in the presence of GAB (a linear regression model of −log_10_p × log_2_fold change through treatment with the control as the independent variable, had a gradient of greater than 1: 1.32, P=0) whereas in GBM63 the gene expression changes were significantly dampened when GAB was added (a linear regression model of −log_10_p × log_2_fold change through treatment with the control as the independent variable, had a gradient of less than 1: 0.016, P=0) (**Fig. 4D**).

These results suggest that modulating GABA signalling affects how GBM cells transcriptionally reprogramme in response to standard treatment, but whether this aids or hinders cell killing depends on the responder subtype.

## Discussion

We have shown that a subset of genes is consistently and significantly dysregulated in locally recurrent IDHwt GBM tumours, following standard treatment, but that the direction of dysregulation is patient specific. This phenomenon classifies patients into Up or Down responders (**Figs. 1D and S1C**). Inspecting the therapy driven transcriptional changes in these subtypes separately suggests different mechanisms of treatment resistance. In Down responders, we observe a predisposition of tumours to become more mesenchymal, both at the bulk level and via an increase in mesenchymal neoplastic cells specifically (**Figs. 2A-B and S2A-B)**. Down responders also exhibit upregulation of proliferation and cell cycling (**Figs. 2C-E and Table S4 and S6**). In keeping with this finding, Wang et al, recently showed an increase in the number of cycling single GBM cells, of the mesenchymal state specifically, at recurrence. The mesenchymal phenotype has been linked with intrinsic resistance to both radiation and TMZ in gliomas[37–39]. This suggests that, in Down responders, there is selection of, or transformation to, more inherently resistant mesenchymal cells during treatment, with subsequent expansion to form a recurrent tumour.

Conversely, in Up responders we observe a higher probability of switching toward a proneural subtype (**Figs. 2A and S2A)**, driven by an increased prevalence of neural- and, especially, oligodendrocyte-progenitor like cancer cells at recurrence (**Figs. 2B and S2B**). We observe a reduction in cell cycling and upregulation of quiescence (**Figs. 2C-E and Table S25 and S27**). Up responders also exhibit a therapy-driven shift towards more differentiated cell states, a unique upregulation of neuronal signalling, and upregulation of signatures observed within the leading edge of GBM tumours which is more enriched with normal brain cells (**Figs. 2C and Table S25 and S27**). GBM cancer cells form synapses with surrounding neurons and glia, creating circuits through which electrical signalling has been shown to promote glioma growth[40–42]. Radiation and TMZ target rapidly diving cells. Our results suggest that, in Up responders, cells evade treatment by reducing proliferation by either quiescing or becoming more differentiated and integrating into normal neural circuits.

Our previous work spanning all types of longitudinal glioma concluded that there are recurrent phenotypes: neuronal, mesenchymal and proliferative, with the former two being specific to IDHwt tumours[8]. Herein we have confirmed these phenotypes but shown that tumours that recur locally following standard treatment can actually be stratified according to phenotype shift at recurrence: Up responders become more neuronal and Down responders become more mesenchymal and proliferative. This suggests that standard treatment is polarising, as also confirmed by Wang et al[9] who initially defined the proneural to mesenchymal axis in patient tissues. Their most recent work shows that tumours become more polarised to either end of this axis during treatment[9].

Cumulatively this has led to our working model of GBM tumour adaption, detailed in **Fig. 4E**. We propose that Down responder tumours (blue dotted arrows) adapt by converting to a more proliferative, mesenchymal state that is able to continue dividing owing to being more intrinsically able to survive standard treatment; Up responders (red dotted arrows) reduce proliferation, thus avoiding death by radiation and TMZ, by converting to a more quiescent, proneural phenotype or becoming more differentiated and able to integrate with normal neural circuits.

Our results raise the possibility that responder subtype stratification will lead to more effective, targeted treatments. To begin testing this we identified *in vitro* models that recapitulate the responder subtypes and tested the effect of adding modulators of GABA signalling alongside radiation and TMZ. We found that this significantly modulated the effect of standard of care but in different directions (**Fig. 4C**). This justifies future work identifying and testing responder subtype-specific targets, such as those identified through our network approach (**Fig. 3C**). The bidirectional nature of subtype transcriptional reprogramming suggests that adjuvant therapies that benefit one subtype could be detrimental to the other, as was borne out in our *in vitro* experiment. This may explain previous failures of clinical trials where retrospective subtyping could elicit whether certain therapies would be more effective post-stratification. However, stratified medicine approaches will only be possible if the responder subtype can be accurately predicted from the primary tumour; work is ongoing in this area.

An alternative therapeutic approach to targeting the biological differences downstream of responder stratification, is to target the mechanism by which responder subtypes derive i.e. the mechanism that underpins reprogramming of the JBSgenes in different directions. Such a target may be universal. Our data implicates JARID2, as the dysregulated genes were identified through commonality of binding this protein in public datasets. JARID2 is an accessory protein to Polycomb Repressive Complex 2 (PRC2), an epigenetic remodelling complex with known roles in brain cell lineage determination and heavily implicated in GBM cell plasticity. Further work investigating a possible role for this epigenetic complex in determining responder subtype is underway.

Our work has identified two responder subtypes in GBM, suggesting that previously identified recurrent phenotypes result from adaption of tumours along a phenotypic axis. The mechanism enabling this adaption, or the biology underpinning the differential responses, yield opportunity for adjuvant treatments to make standard of care more effective for patients with this deadly disease.

## Methods

Data analyses were performed in R[43] and Python 3 with plots were made using ggpubr where detailed[44].

### Sample collection and processing

Longitudinal GBM samples were acquired from The Walton Centre, Lancashire Teaching Hospitals and Leeds Teaching Hospitals National Health Service (NHS) Foundation Trusts via the Brain Tumour Northwest Tissue Banks and the Leeds Neuropathology Research Tissue Bank. In addition, tissue samples were obtained from Cambridge University Hospitals NHS Foundation Trust as part of the UK Brain Archive Information Network (BRAIN UK)[45]. Samples were processed as previously described[46]. Briefly, formalin-fixed paraffin-embedded (FFPE) blocks were sectioned and the first and last section were H&E-stained and underwent neuropathologist review to identify areas of >60% tumour. Regions of overlap were macrodissected from the intervening sections and RNA was extracted using AllPrep DNA/RNA FFPE Kit (catalogue #80234) from Qiagen (UK).

### RNAseq data acquisition and processing

All RNA extracted in house underwent rRNA depletion using the NEBNext rRNA Depletion Kit (Human/Mouse/Rat) and then strand-directional, whole transcriptome library preparation using NEBNext Ultra™ II Directional RNA Library Prep Kit for Illumina®, both from New England Biolabs (UK). Libraries were sequenced on Illumina next-generation sequencers as 100bp paired end reads. Raw RNA data was acquired from several published studies following the negotiation of Data Transfer Agreements, where necessary[2, 47–50] (See **Table S1**). All Discovery cohort, and *in vitro*, FASTQ data were trimmed of low-quality bases, phred threshold=20, and adapters via Trim Galore v0.4.3, wrapping Cutadapt v1.8.3[51]. Trimmed reads were quality checked using FASTQC[52] and then aligned to human reference genome GRCh38.13 using STAR v020201 in two-pass mode with a maximum of 5 multireads[53]. Gene and transcript count and gene expression was quantified via CuffQuantv2.2.1 taking directional specifics of the library as input, using probabilistic weighting of multireads and quantifying against the GENCODEv27 human genome annotation with haplotypes and scaffolds included[54, 55]. Sequencing metrics are given in **Table S2**. Validation cohort data (**Table S1**) was acquired as pre-processed transcript counts and trasncripts per million (TPM) via the GLASS portal at https://www.synapse.org/glass[8]. Transcript expression was converted to gene-level or transcription start site (TSS)-level data by summing isoform expression. Genes were filtered to keep only those that were expressed above the lower quartile of non-zero gene expression in at least 25% of either primary or recurrent samples.

### Differential expression analysis

Differential expression (DE) analysis was performed using DESeq2 using a paired design[56]. Functional enrichment was assessed via WebGestalt (v2019)[57] with gene set size<1000 on DE genes with FDR<0.1. The DE enrichment diagram was created using enrichplot R package[58]. The statistical significance of the overlap between differentially expressed genes from different cohorts was inferred using the hypergeometric method as implemented in http://nemates.org/MA/progs/overlap_stats.html. DE was also run separately on Up and Down responders with results given in tables S22-23. To evaluate differences in the directions of dysregulation of gene sets between responder types, 2×2 contingency Chi squared tests were performed on the numbers of genes that significantly increased or decreased through treatment in Up and Down responders, in addition to 2×3 Chi squared tests on the numbers that increased, decreased or remained stable. The raw p-values as well as adjusted FDRs are given alongside GSEA results in **Tables S25-27**.

### Gene set enrichment analysis

We developed a novel gene set file for use in GSEA using the Gene Transcription Regulation Database (GTRD v19.10)^[59]^. A gene was assigned to a DNA-binding factor’s gene set if its promoter (transcription start site from gencodev27 ±1kbp, or ±2 or 5kbp where specifically stated) contained a binding site for that factor in ≥2 independent ChIPseq experiments. We first performed pre-ranked GSEA[60], per patient, ordering genes by the magnitude of fold change in expression log_2_(|recurrent expression +0.01/primary expression +0.01|) in classical mode. To indicate directionality of dysregulation we then ranked genes by fold change, referred to throughout as Log_2_FC i.e. using log_2_(recurrent expression+0.01/primary expression +0.01), and weighted by magnitude^[61]^. Cell lines were processed the same as the above, with arbitrarily paired untreated and treated replicates representing primary and recurrent samples. When running GSEA on Up and Down responders separately, genes were ranked and weighted based on the log2 of their significance in differential expression from primary to recurrent. All GSEA runs were set for 1,000 permutations and to allow gene set sizes of up to 50,000. Results are given in Tables S3-18. Heatmaps were created using ComplexHeatmaps R package[62]. Gene set network diagrams were created using enrichplot R package[58].

### Tumour subtyping and cell deconvolution

GBM subtype calls were performed using GlioVis[63] with the 3-way assignment used. Sankey plots were created with SankeyMATIC https://sankeymatic.com. IDHwt GBM tumour specific cell deconvolution was done using GBMDeconvoluteR using the default gene markers with non-tumour intrinsic genes filtered out[10, 26].

### Survival analysis

A multivariate linear regression was performed to assess the relation between progression free and overall survival (months) and the explanatory variables: Age (years), batch corrected MGMT expression (transcripts per million) in the primary tumour and JARID2 gene set normalised enrichment score. Data were checked for multicollinearity with the Belsley-Kuh-Welsch technique. Heteroskedasticity and normality of residuals were assessed respectively by the Breusch-Pagan test and the Shapiro-Wilk test. A p-value < 0.05 was considered statistically significant. Statistical analysis was performed with the online application EasyMedStat (version 3.16; www.easymedstat.com).

### Gene correlation networks

Log2FC values were used to build correlation networks for Up and Down responders separately with Parsimonious Gene Correlation Network Analysis (PGCNAv2)[64] using the Ledenalg algorithm for community detection. Default parameters were used except for setting “ –pgcna_f 1, --pgcna_n 1000 “ to ensure that all genes were used in network construction and that the clustering algorithm is run 1000 times to enable the optimal network to be assessed, and selected, according to the Scaled Cluster Enrichment Score (SCES). An integrated value of influence (IVI) score was assigned to each gene using the influential R package[65].

### Preparation of stock solutions

Temozolomide (Merck, T577-100MG) was resuspended to 50 mM in dimethyl sulfoxide (DMSO) and stored at −20°C. Gaboxadol hydrochloride (Merck, T101-500MG) and Guanidinoethyl sulfonate (Cayman Chemical, 17572-500mg-CAY) were both resuspended in high purity water to 100 mM and stored at −20oC. New stocks of GAB and GES were made every 6 months.

### Cell line acquisition and culture

The A172 established GBM cell line was acquired ATCC (CRL-1620). A172 cells were maintained in DMEM (Merck, D6429) and 10% fetal bovine serum (FBS). GBM58 and GBM63, derived in Leeds, were cultured in NB media (ThermoFisher Scientific, 10888022) supplemented with 40 ng/mL recombinant human EGF (Peprotech, 236-EG-200) 40ng/mL recombinant human FGF (R&D systems, 100-18B-100), 0.5 × B27 serum free supplement (ThermoFisher Scientific, 17504044) 0.5 × N2 supplement (ThermoFisher Scientific, 17502048). GBM58 and GBM63 were cultured in flasks coated with 10 μg/mL ornithine (Sigma, P3655-50MG) and 2 μg/mL laminin (Sigma, L2020-1MG). Cells were maintained at 5% CO2 at 37°C and passaged when at 80% confluency. On passage of the cells, they were washed with 5mL Dulbecco’s Phosphate Buffered Saline (PBS) before addition of Trypsin-EDTA solution at 1 mL/75cm2 flask. Cells were placed in the incubator until detached before being collected in the appropriate media and centrifuged for 5 minutes at 300g before media being removed. Cells were resuspended in medium and split at appropriate confluency. For plate coating, poly-L-ornithine stocks were diluted to 10 μg/mL in TC grade water. 10 mL working solution was added to each T75 flask. After one hour at room temperature, the solution was removed, and flasks rinsed with TC grade water. Laminin stocks were diluted to 2 μg/mL in PBSA. 10 mL working solution was added to each T75 flask. Flasks and plates were wrapped in parafilm and left at room temperature overnight before storing at −20°C.

### Spheroid culture

For 3D culture, cells were trypsinised and resuspended at 1.5×104 cells per mL in normal culture medium supplemented with 10 mM taurine and 200 μL plated into each well of a 96 -Well Clear Round Bottom Ultra-Low-Attachment Microplate (Scientific Laboratories Supplies, 7007). Any empty wells were filled with 200 μL PBS to avoid evaporation. Spheroids were imaged immediately before treatment and one week post treatment using the Confocal Nikon AR1 and medium changed every three days by removing 100 μL of medium and replacing this with 100 μL fresh medium.

### Treating with temozolomide and irradiation

At 5 days post-seeding, 100 μL of medium was removed from each spheroid and replaced with 100 μL medium containing TMZ diluted from the 50 mM stock solutions to 60 μM, giving a final concentration when added to spheroids of 30 μM. One hour after TMZ addition, cells were irradiated using a RadSource RS-2000 X-ray irradiator with 2Gy.

### Inhibitor and standard treatment

At 5 days post-seeding, 120 μL of media was removed from each spheroid. For vehicle control spheroids 20 μL dH2O was added. 20 μL of 100 mM GES or GAB was added to remaining spheroids. Following this, untreated spheroids received 100 μL of media containing 10 mM taurine, and the spheroids to be treated with standard treatment received 100 μL media containing 10 mM taurine and 60 μM TMZ to give a final concentration of 30 μM TMZ. Treated plates were subjected to irradiation treatment using a RadSource RS-2000 X-ray irradiator with 2Gy one hour after TMZ treatment. Experimental end point assays of imaging and Cell-Titre glo 3D cell viability assay were performed 7-days post treatment.

### Spheroid imaging and growth curves

To measure spheroid growth, a bespoke automated plate-imaging and analysis programme was developed using the Confocal Laser Scanning Microscope-Nikon A1R. Area (μm^2^) would be the measurement used to represent spheroid size. Data was analysed using SpheroidAnalyseR[66] which uses a pre-set threshold to remove obvious outliers for example empty wells, and then removes further statistical outliers using a Robust-Z-Score of +/− 1.96. Linear mixed effects models were built using R package lme4 v1.1.28[67]. The significance of the interaction between the effect of treatment and inhibitors on spheroid size was calculated, separately for each cell line, via Anova between the null model [Size ~ Treat + Inhibitor+ (1+Treat|Subject)] and that which included the interaction term [Size ~ Treat*Inhibitor+ (1+Treat|Subject)]. Effect sizes and significances were then calculated for the interaction models fitted by maximum likelihood.

### CellTitre-Glo 3D cell viability assay

Spheroids to be analysed via CellTitre-Glo 3D cell viability assay were removed from the spheroid plate in a total volume of 100 μL and transferred to 96-well white opaque edged plates (Grenier Bio-one Ltd) and left at room temperature for 30 minutes. CellTitre-Glo reagent was settled at room temperature before 20 μL was added to each well. Plates were placed on a plate shaker for 30 minutes at room temperature before the luminescence read on a Cytation 5 Imaging Plate Reader (BioTek).

### RNA from spheroids for sequencing

Around 50 spheroids per condition were collected in a 15 mL centrifuge tube and centrifuged at 800 rpm for 5 minutes. Media was aspirated and spheroids washed in PBS and centrifuged at 800 rpm for 5 minutes. PBS was removed and spheroids washed in PBS and centrifuged at 800 rpm for 5 minutes. PBS was removed and 600 μL of Qiazol added from Qiagen Lipid Tissue Mini Kit. Spheroids in Qiazol were frozen at −80°C for 24 hours before being defrosted. Once defrosted, Qiagen Lipid Tissue Mini Kit (Qiagen, 70804) was used to extract RNA as per the manufacturer’s instructions. RNA was quantified using Nanodrop spectrophotometer (ThermoFisher) assessment before being stored at −80°C. RNA was sequenced and analysed as per that from patient samples.

## Declarations

### Ethical Approval

All samples were from patients that provided informed, written consent. The use of consented samples was following favourable opinion of a project-specific application by NHS Health Research Authority: South Central - Oxford A Research Ethics Committee (reference 13/SC/0509).

### Data Availability

The raw data created for this study, in the form of sequenced reads, are available from the European Genome-phenome Archive repository (https://www.ebi.ac.uk/ega/home), accession EGAD00001009806 upon completion of a Data Transfer Agreement with the University of Leeds. Data taken from published studies is available as described in the original publications listed in **Table S1**.

### Code Availability

Code is available on GitHub via https://github.com/GliomaGenomics/GBM_TF_analysis

### Competing Interests

RGWV is a consultant for NeuroTrials, Inc, and Stellanova Therapeutics. AD is director of AD Bioinformatics Ltd

### Funding

This work was supported by grants from UK Research and Innovation [MR/T020504/1 to LFS], Leeds Hospitals Charity [9R11/14-11 to LFS], Brain Research UK [PhD Studentship to LFS for RB], the Medical Research Council [DiMeN PhD Studentship to LFS for NA] and Health Data Research UK, an initiative funded by UK Research and Innovation, Department of Health and Social Care (England) and the devolved administrations, and leading medical research charities [MSc Studentship for CH]. The Brain Tumour Northwest Tissue Banks at The Walton Centre and Royal Preston Hospital are funded by Sidney Driscol Neuroscience Foundation. The Leeds Neuropathology Research Tissue Bank and SP, are funded by OSCAR’s Paediatric Brain Tumour Charity and Yorkshire’s Brain Tumour Charity [Joint Infrastructure Funding to LFS]. The UK Brain Archive Information Network (BRAIN UK)[45] is supported by Brain Tumour Research, the British Neuropathological Society, the Medical Research Council and brainstrust.

### Author contributions

LFS conceived and designed the study. MDJ and AB supplied samples; KS and AB supplied clinical metadata. AC and AI performed neuropathology review and annotation. NR, AFB, MF and SP processed patient samples and extracted RNA. SS supervised AFB. GT, NA, SA, CH and LFS analysed data, with input from FSV, AD and RGWV. MAC, LC, DW and LFS supervised NA. AG and LFS supervised SA. RB completed all *in vitro* experiments with input from MF. JP, EW, CJ and LFS supervised RB with input from ABR. LFS wrote the manuscript. All authors read and approved the final manuscript.

