## Supplemental Figures for "IDHwt glioblastomas can be stratified by their transcriptional response to standard treatment, with implications for targeted therapy"

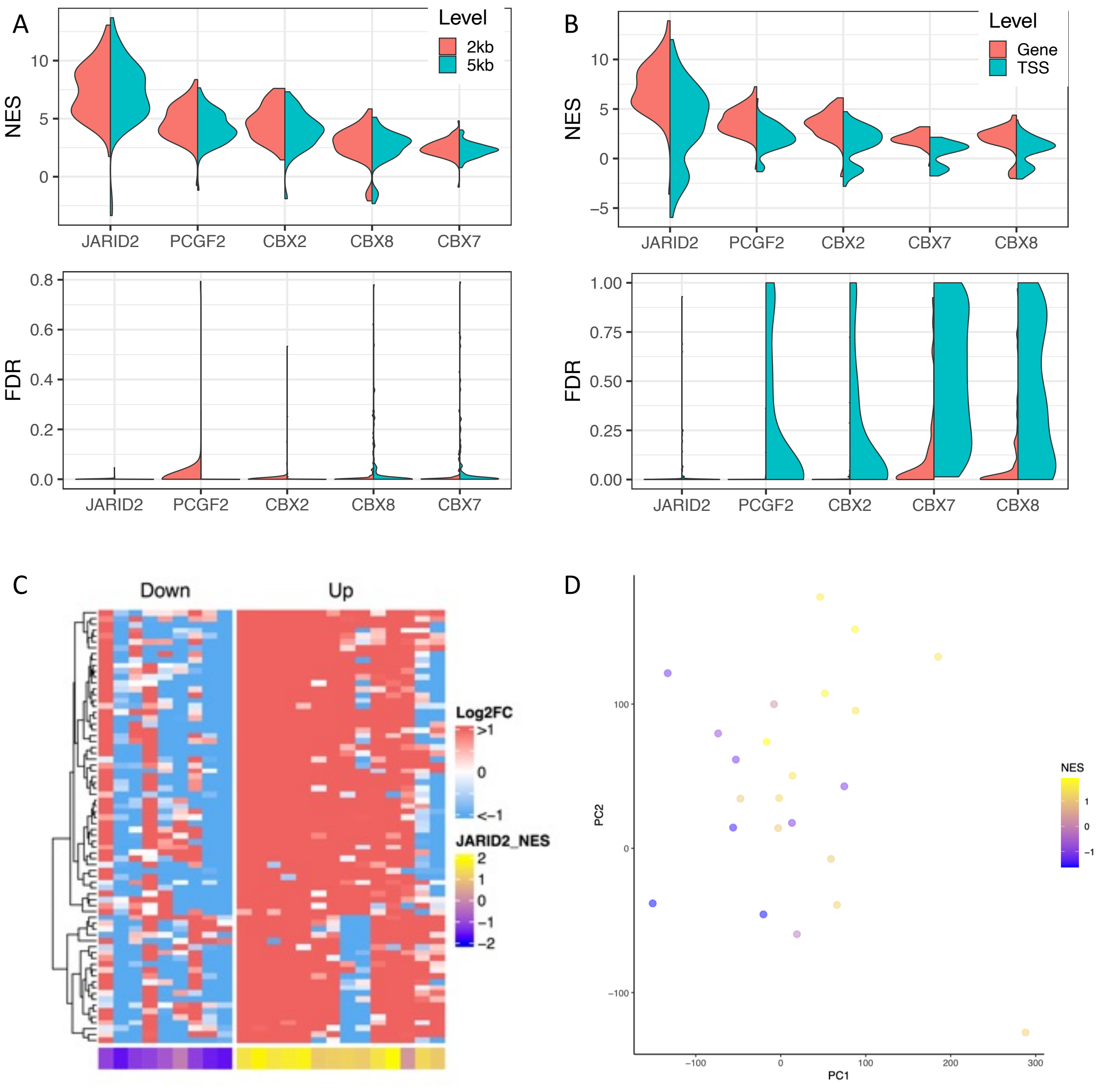


**Figure S1. A**. The distribution of per-patient normalised enrichment scores (NES, top plot) and false discovery rates (FDR, bottom plot) for the top-scoring promoter-binding factors associated with gene expression changes between recurrent vs primary GBMs, in the Discovery cohort, when the promoter definition is changed from 1kb to 2kb (pink) or 5kb (teal) either side of the transcription start site (TSS). **B.** As panel A except results are for the TSS $\pm$1kb definition of a promoter but using expression at the level of individuals genes (salmon pink) or resulting from each unique TSS (teal). **C.** A heatmap of the fold change in expression between recurrent vs primary GBMs samples for each patient (columns) for the JARID2 binding sites genes (JBSgenes) that were in the leading edge of the GSEA results in more than 50% of patients (LE50 genes, rows) in the Validation cohort. **D**. Validation cohort patients are plotted, coloured by JBSgenes NES, according to principal components 1 (PC1) and 2 (PC2) of their whole transcriptome fold change in expression between their recurrent vs primary GBM.

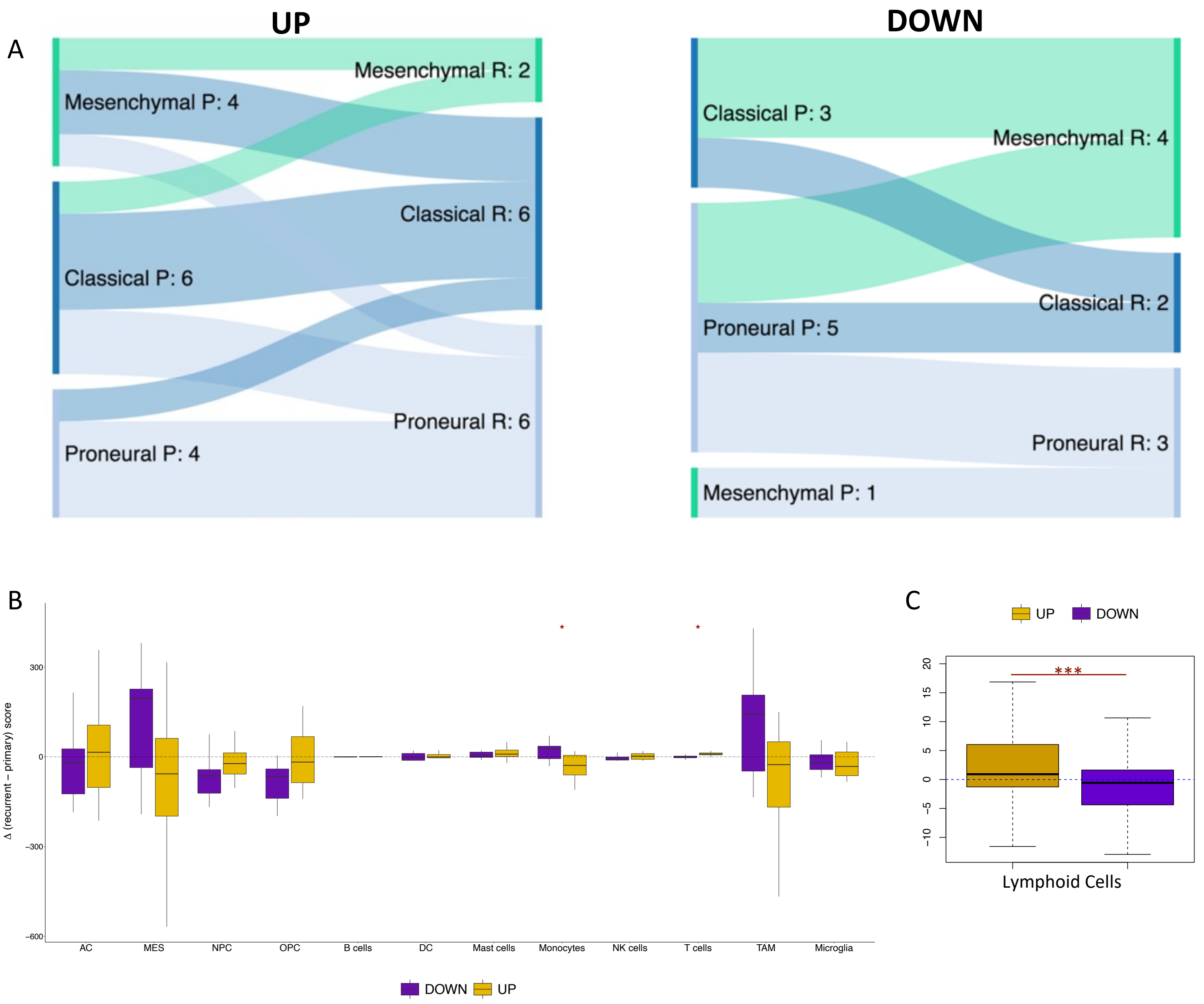


**Figure S2. A.** Sankey plots showing the prevalence of subtype switching from primary (P) to recurrent (R) GBM in the Up responders (left) and Down responders (right) in the Validation cohort. **B.** The distributions of change in cell type score, assigned per sample by GBMdeconvoluteR to indicate the prevalence of that cell type in the tumour, between primary and matched recurrent GBMs in Down (purple) and Up (gold) responders in the Validation cohort. A dotted line indicates no change. The median is noted by a black horizontal line. Significance is denoted by asterisks: *: $p<0.05$; **: $p<0.01$; ***: $p<0.001$; ****: $p<0.0001$. Neoplastic GBM cells are on the left of the plot: AC= astrocyte like; MES= mesenchymal like; NPC= neural progenitor like; OPC= oligodendrocyte progenitor like. **C.** As per B but after the Discovery and Validation cohorts were combined and immune cells relabelled as either lymphoid (T cells, B cells, NK cells) or myeloid (DC cells, mast cells, monocytes, microglia and tumour-associated macrophages).

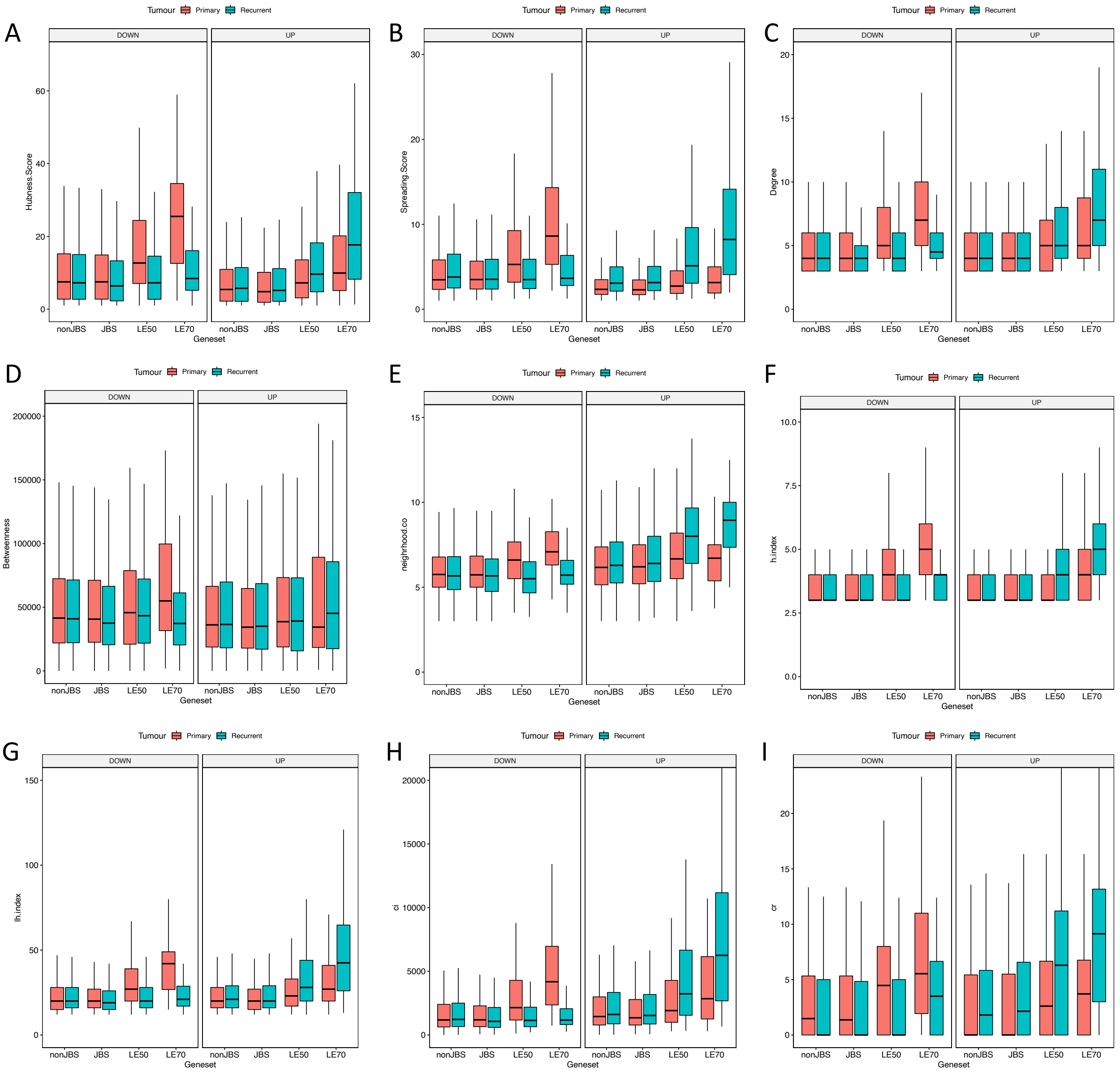


**Figure S3.** Network metric distributions for different gene sets, calculated from correlation networks that were built from gene expression data in primary (pink) or recurrent (teal) tumours in Down (left panel) or Up (right panel) responders, separately. The network metrics are **A.** Hubness. **B.** Spreading. **C.** Degree. **D.** Betweenness **E.** Neighbourhood connectivity **F.** H-index (h.index) **G.** Local H-index (lh.index) **H.** Collective influence (ci) **I.** Cluster rank (cr). nonJBS: genes not in the JARID2 gene set; JBS: genes in the JARID2 geneset but excluding those in the leading edge of at least 50% of patients (LE50 genes); LE50: genes in the LE50 geneset but excluding those in the LE70 geneset; LE70: genes in the leading edge of at least 70% of patients.
